# A prefrontal to lateral entorhinal pathway disrupts memory

**DOI:** 10.1101/2022.02.21.480888

**Authors:** Erin Hisey, Scott H. Soderling

## Abstract

While the neural circuits underlying memory encoding, storage, and retrieval are well characterized, the circuits that act to disrupt memory are enigmatic. Here we find that silencing a projection from the prefrontal cortex to the lateral entorhinal cortex surprisingly improves spatial working memory and contextual memory. We then found that the same cell type shows increased activity during errors in a spatial working memory test. Finally we found that optogenetic activation of the same activity patterns can disrupt working memory performance. By using a combination of intersectional genetic and in vivo imaging techniques we advance evidence that a novel prefrontal-entorhinal pathway critically participates in memory disruption.

## Introduction

Memory is an essential and diverse neuronal faculty that operates at multiple time scales (i.e., long-term and short-term or working) and information types (i.e., spatial, non-spatial, declarative, and non-declarative). As such, multiple circuit and signaling mechanisms underlie these varied information storage processes. The prefrontal cortex (PFC) and hippocampus play crucial roles in working memory and long-term memory storage, and contextual and spatial memory, respectively [1, 2]. Though these two memory centers are critical for multiple forms of memory and intensively studied across humans and rodent models, the circuits that connect the two are less well understood. The lateral entorhinal cortex (LEC) receives dense reciprocal projections to and from the PFC as well as the hippocampus, serving as a critical hub between these two areas [3]. LEC is known to encode complex associations between contexts and objects in rodents and is critical for long-term memory in humans [4–7]. The projection from the PFC to the LEC, in particular, is poised to regulate information flow to the hippocampus while the cell bodies themselves are located throughout prelimbic, infralimbic, and orbitalfrontal regions of PFC (Supp. Fig. 1). However, the functional role of these PFC cells projecting to LEC (PFC-LEC cells) in memory is unknown.

As both the PFC and LEC are implicated in multiple forms of memory, we hypothesized that PFC-LEC cells would be necessary to facilitate forms of memory both PFC and LEC play a role in. To our surprise, rather than facilitate memory, we found that PFC-LEC cells instead appear to disrupt memory.

### Inactivation of PFC-LEC cells improves working and contextual memory

To examine whether PFC-LEC cells regulate forms of memory the PFC and LEC have been previously implicated in, we chose two tasks, Y maze and novel object-in-context, given their demonstrated dependence on the prefrontal cortex, and the LEC, respectively, in order to test if PFC-LEC cells play a role in both, either, or none of the functions previously attributed to the PFC and LEC. We selectively silenced PFC-LEC cells by injecting a virally encoded Cre-dependent tetanus toxin [8] into the PFC and a retrogradely transported Cre [9] into the LEC of adult mice (Fig.1a). After allowing 4 weeks for viral expression, experimental mice were first analyzed in the Y maze. In this unrewarded task, working memory is thought to be intact if mice spontaneously alternate through each arm of the maze and avoid subsequently revisiting arms just visited. To our surprise, the experimental mice improved in their ability to navigate through the Y maze compared to control siblings injected with Cre-dependent tetanus toxin only. The experimental group made significantly more alternations through the maze (Fig.1b), and significantly fewer direct revisits (Fig.1c) compared to controls. The increase in alternations was not simply due to hyperactivity in the experimental group as the PFC-LEC silenced mice did not differ from controls in the distance they traveled (Fig. 1d).

**Figure 1:**
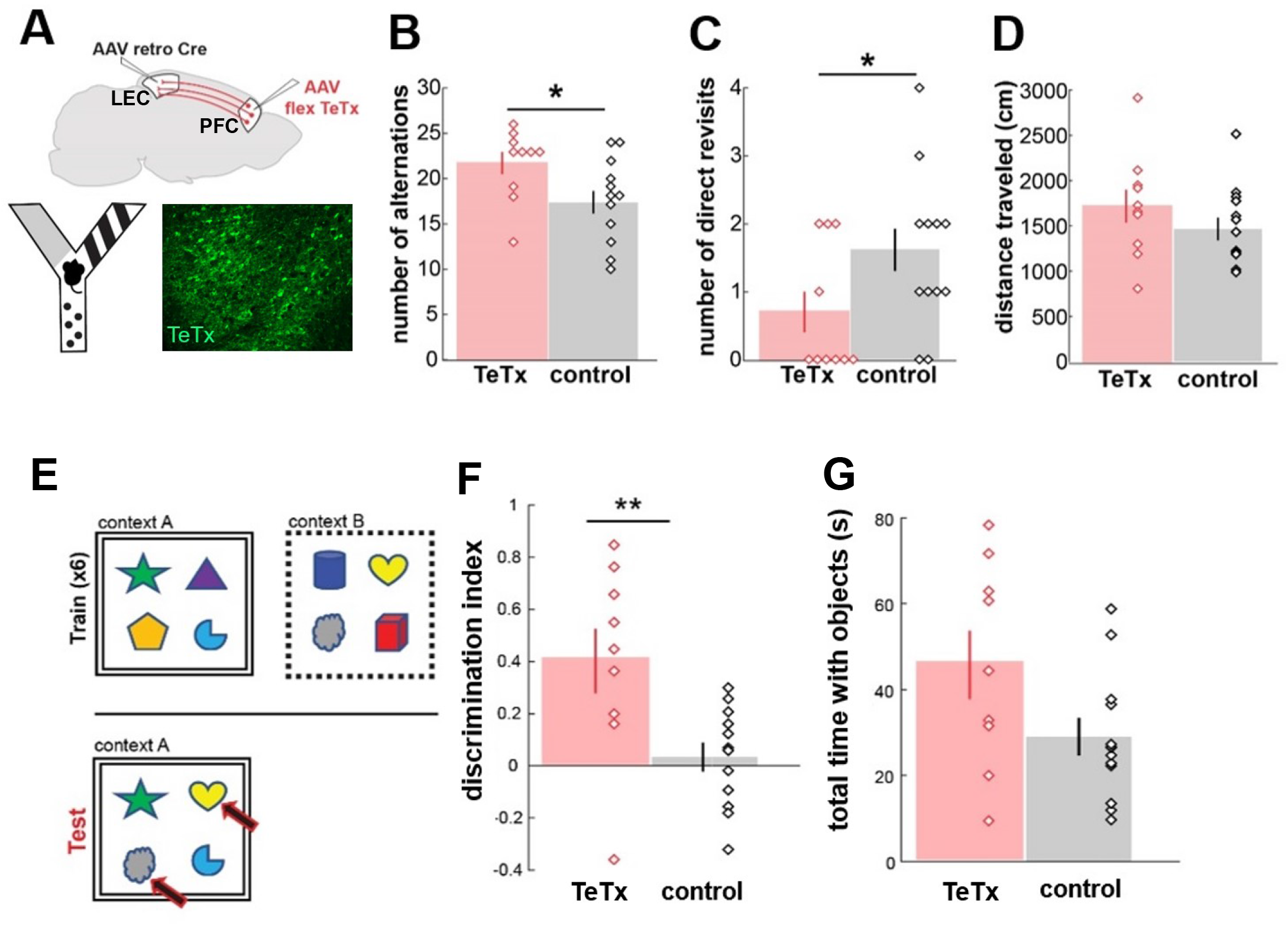
Intersectional genetic silencing improves working memory and contextual memory. A) Schematic showing viral strategy, schematic of Y maze as a test for working memory, and representative TeTx-YFP label in PFC. Note that mice were placed in the maze for 5 minutes. B) Number of alternations in experimental (TeTx, n = 10) and control animals (control, n = 13) (two-tailed t-test p-value = 0.03). C) Number of direct revisits (two-tailed t-test p-value = 0.05). D) Total distance traveled during testing. E) Schematic of object-in-context task. F) Discrimination index in experimental (TeTx, n = 9) and control animals (control, n = 13) (two-tailed t-test p-value = 0.008). G) Total time with objects.

We then wanted to determine if PFC-LEC cells, role in working memory suppression extends to other forms of memory that are implicated in both PFC and LEC function. Both the PFC and LEC are critical to contextual memory, given that lesioning the PFC and the LEC in rats impairs object-contextual memory while leaving basic object recognition intact [10–12]. We tested the same cohort of PFC-LEC inactivated mice in an object-context task adapted for use in mice [13] (Fig. 1e). Control mice discriminate relatively poorly between familiar objects in familiar contexts and familiar objects in novel contexts. In contrast, the experimental mice discriminated significantly better between novel object-context pairings, spending more time with familiar objects in novel contexts than with objects that remained in the same context as in training (Fig. 1f). The improved discrimination cannot be accounted for by an increase in object exploration during testing as the experimental and control mice did not significantly differ in total time spent with objects (Fig 1g). Differences in object preference and exploration during training can also not account for the improved discrimination seen in testing. Experimental mice showed no differences to controls in initial preference for objects or arena exploration during training (Supp. Fig. 2). Taken together, the improvement of both working memory and contextual memory tasks when PFC-LEC cells are silenced suggests that, rather than facilitate memory, PFC-LEC cells may interfere with multiple forms of memory.

### FC-LEC cells show differential responses in a working memory task

To probe how PFC-LEC cells might interfere with working memory in particular, we drove the expression of a fluorescent calcium indicator (GCaMP6f) [14] specifically in PFC-LEC cells and implanted mice with a GRIN lens over PFC (Fig. 2a). We then coupled the mice to a miniature microscope and collected calcium activity from PFC-LEC cells while mice spontaneously explored the Y maze (behavior in Y maze, Fig. 2g, Supp. Fig. 3). We first examined the event rate of PFC-LEC cells at different locations in the Y maze to determine if activity was differentially modulated at the center of the Y maze, the maze region in which working memory is thought to be engaged. Overall, PFC-LEC cells trend towards higher event rates while in the center of the Y maze than the arms of the Y maze with a slope significantly lower than one, with one being expected if center and arm rates were identical (Fig. 2b, slope of center rate versus arm rate = 0.614, 95% confidence intervals: 0.536 to 0.692). We next wanted to determine if cells that showed selectivity for the center of the maze over the arms showed differences in event rates when entering and exiting the center of the Y maze. This analysis was important as entry into the center of the Y maze is the portion of the task most heavily dependent on working memory, as the mouse retains the memory of which arm of the maze most recently visited before choosing the next arm to visit [15, 16]. Selectivity was defined as (center rate - arm rate) / (center rate + arm rate), with center selective cells being those with a selectivity greater than 0 and arm selective cells being those with a selectivity less than 0. In the first second of entering the center of the maze, center selective cells showed a significant increase in activity (Fig. 2c) that was not seen in arm selective cells upon entering the center (Fig. 2e) or upon entering arms of the maze (Fig. 2f). No significant change in activity was observed upon exit of the center in either center-selective (Fig. 2d) or arm selective cells (Fig. 2f). This unique and significant increase in activity upon center entry in center-selective cells suggested that PFC-LEC cells were indeed preferentially active at a time in which working memory is engaged, positioning them at an optimal time at which to interfere with memory processing. We then hypothesized that if PFC-LEC cells actively interfere with memory, they should show increased activity in the maze center before an incorrect visit to an arm. We compared the activity of PFC-LEC cells in the center of the maze during correct alternations to their activity during relatively rare mistakes (“direct revisits”) (Figure 2G). While we saw little modulation of activity in the center of the maze between non-consecutive visits to the arms of the maze (“alternation”, n = 39 alternations from 6 mice, Fig. 2i), we found a significant increase in events during the center visit before an incorrect choice compared to the previous visit to the center (“direct revisit”, n = 9 direct revisits from 4 mice, Fig. 2h). These data are consistent with PFC-LEC cells playing an instructive role in working memory. In particular, the elevated activity preceding incorrect choices suggested the activity of this circuit could potentially represent active interference with working memory processing.

**Figure 2:**
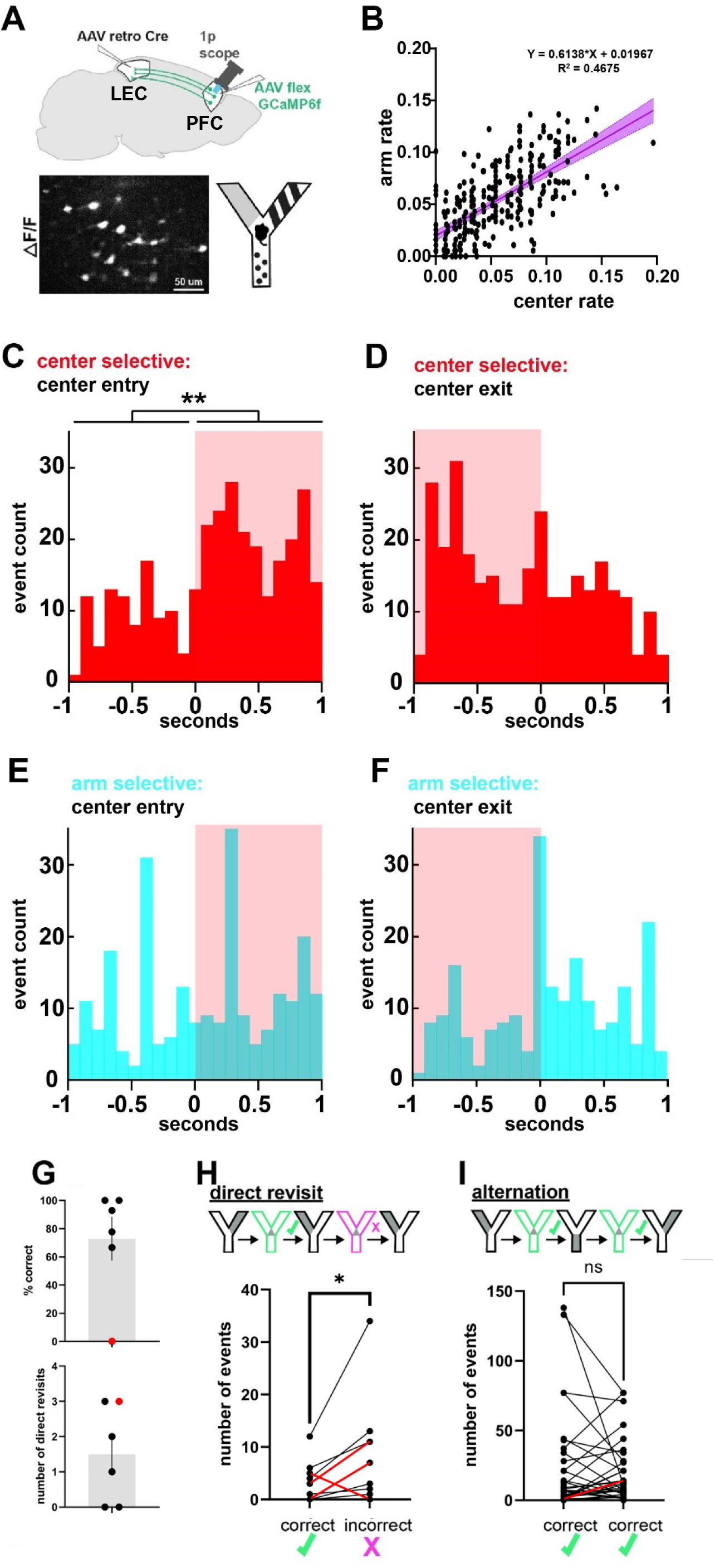
FC-LEC cells show differential responses in a working memory task. A) Schematic of viral injection and lens implant. B) Event rate in center versus event rate in the arms for each cell (n =276 cells from 6 mice; slope = 0.614, 95% confidence intervals: 0.536 to 0.692, R^2^ = 0.4675) C) Histogram of all events of center selective cells (n = 135 center selective cells from 6 mice) aligned to center entry. Shaded area indicates events when the mouse is in the center of the maze (Kolmogorov-Smirnoff test between summed activity distribution 1 second before and during time in center, p = 0.004). D) Histogram of all events of center selective cells aligned to center exit in all mice (Kolmogorov-Smirnoff test between activity distribution during 1 second in center and 1 second after center exit, p = 0.209). E) Histogram of all events of arm-selective cells (n = 141 arm selective cells from 6 mice) aligned to center entry (Kolmogorov-Smirnoff test between summed activity distribution 1 second before and during time in center, p = 0.697). F) Histogram of all events of arm selective cells in all mice aligned to center exit (Kolmogorov-Smirnoff test between summed activity distribution during 1 second in center and 1 second after center exit, p = 0.208). G) Behavior in Y maze during Inscopix imaging. Top, percent correct ([complete alternations - direct revisits]/complete alternations)*100). Bottom, number of direct revisits. Mouse with poor performance is indicated in red. H) Center events during direct revisits. Number of center events before correct arm choice followed by number of center events before incorrect arm choice (n = 9 direct revisits from 4 mice, one-tailed paired t-test between each revisits, p = 0.0226 ratio paired t-test between total events for each mouse, p = 0.013). Red lines indicate events for mouse with poor performance. I) Center events during correct alternation, determined by the number of center events before correct arm choice followed by number of center events before another correct arm choice. Red line indicates events for mouse with poor performance.

### Temporally specific activation of FC-LEC cells disrupts working memory

We next sought to determine if the increased activity upon center entry, observed in our one-photon imaging, is critical for PFC-LEC cells to interfere with working memory. We hypothesized that increased activation of PFC-LEC cells on center entry would impair working memory performance in the Y maze. To test this possibility, we injected virally encoded Cre-dependent channelrhodopsin [17] in the PFC and a retrogradely transported Cre in LEC. Fiberoptic cannulas were implanted near the midline of the PFC. Labeled PFC-LEC neurons were exposed to triggered light stimulation for one second at 10 Hz upon entry to the center of the Y maze (Fig. 3a). Consistent with our hypothesis, light stimulation decreased the number of alternations in experimental mice compared to implanted control siblings injected with GFP (Fig.3b). The number of direct revisits also trended towards an increase in the experimental group compared to controls but not significantly so (Fig. 3c). Temporally specific activation of PFC-LEC cells did not cause a change in locomotor activity as distance traveled did not significantly differ between experimental and control groups (Fig. 3d). Additionally, the reduction in the number of alternations was not mediated by increased time spent in the center of the maze, suggesting that stimulation of PFC-LEC cells does not provide an appetitive signal that drives the mice to spend more time in areas where they receive light stimulation (Fig. 3e). That specific activation of PFC-LEC cells during a time window in which they were found to be endogenously active results in impaired working memory performance, complements our finding that silencing the same cells improves working memory. This data further suggests that PFC-LEC cells are engaged in a mechanism that interferes with, rather than promotes, memory.

**Figure 3:**
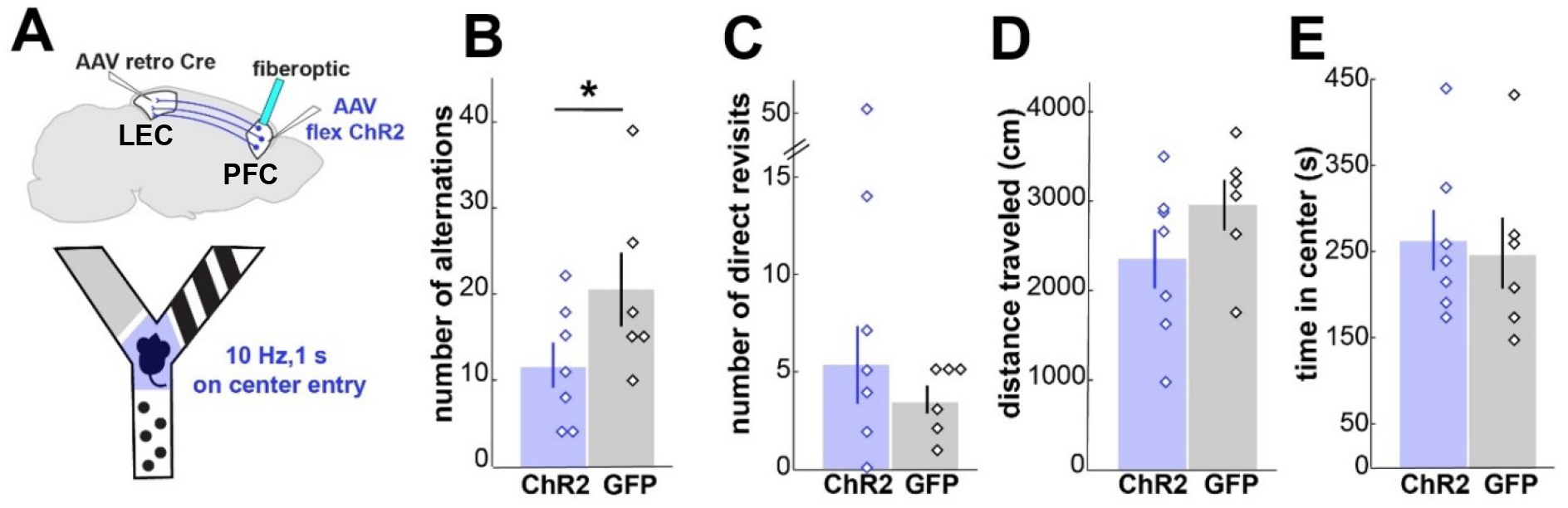
Temporally specific activation of PFC-LEC cells decreases the number of alternations. A) Schematic of viral strategy and behavioral paradigm. Note that mice were placed in the Y maze for 10 minutes. B) Total number of alternations in the Y maze in fiberoptic implanted animals with channelrhodopsin (ChR2; n = 7) or GFP (GFP; n = 6) expressed in PFC-LEC cells (one-tailed t-test p-value = 0.049). C) Total number of direct revisits. D) Total distance traveled. E) Total time spent in the center of the maze in seconds.

## Conclusion

Here we have demonstrated that silencing of PFC-LEC cells surprisingly improves both working memory and contextual memory. We then show that PFC-LEC cells selectively increase activity upon center entry before an incorrect choice in the Y maze and that activation of these cells specifically in this time window is sufficient to disrupt working memory. The Y maze task used to assay working memory also provides contextual information to the animal via different patterns of the walls of each arm and can be seen as a test of contextual memory as well as working memory so we believe that our optogenetic and imaging results may also extend to contextual memory, though further experiments addressing contextual memory specifically are warranted.

Why might disruption of memory play an essential role in behavior? Connections between the prefrontal cortex and hippocampus have been hypothesized to play a critical role in inhibitory control over memory, facilitating what is termed “retrieval suppression” in humans [24]. As PFC-LEC cells provide monosynaptic excitatory drive to the LEC, they may support top-down suppression of hippocampal retrieval. Another possible explanation of the improvement and impairment in working memory we see when PFC-LEC cells are inactivated and activated, respectively, is that they facilitate active forgetting, a process by which synapses that store memory are remodeled such that the previous memory is no longer available [25]. To distinguish between inhibitory control, where the memory is still present but suppressed, and active forgetting, where the memory trace is actively eroded, future studies would need to use a task design in which memory is tested during an acute manipulation and then after the manipulation to determine if the memory is still present.

---

**Supplementary figure 1:**
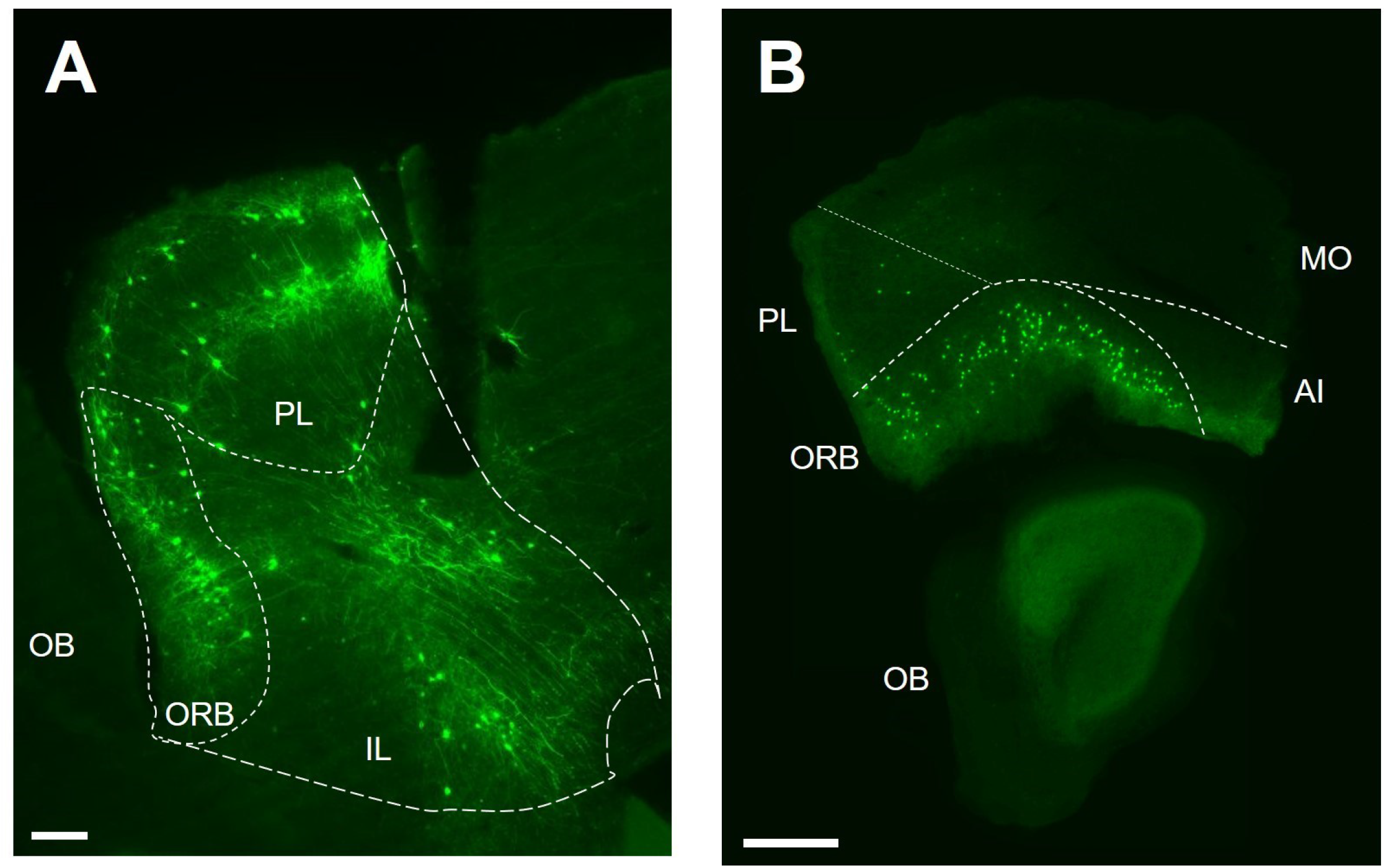
Anatomical location of PFC-LEC cells within the prefrontal cortex. A) GFP-labeled PFC-LEC cells shown in sagittal section through PFC (~0.3 mm M/L). Scale bar, 200 microns. B) GFP-labeled PFC-LEC cells shown in anterior (~2.5 mm A/P) coronal section of PFC. Scale bar, 500 microns.

**Supplementary figure 2:**
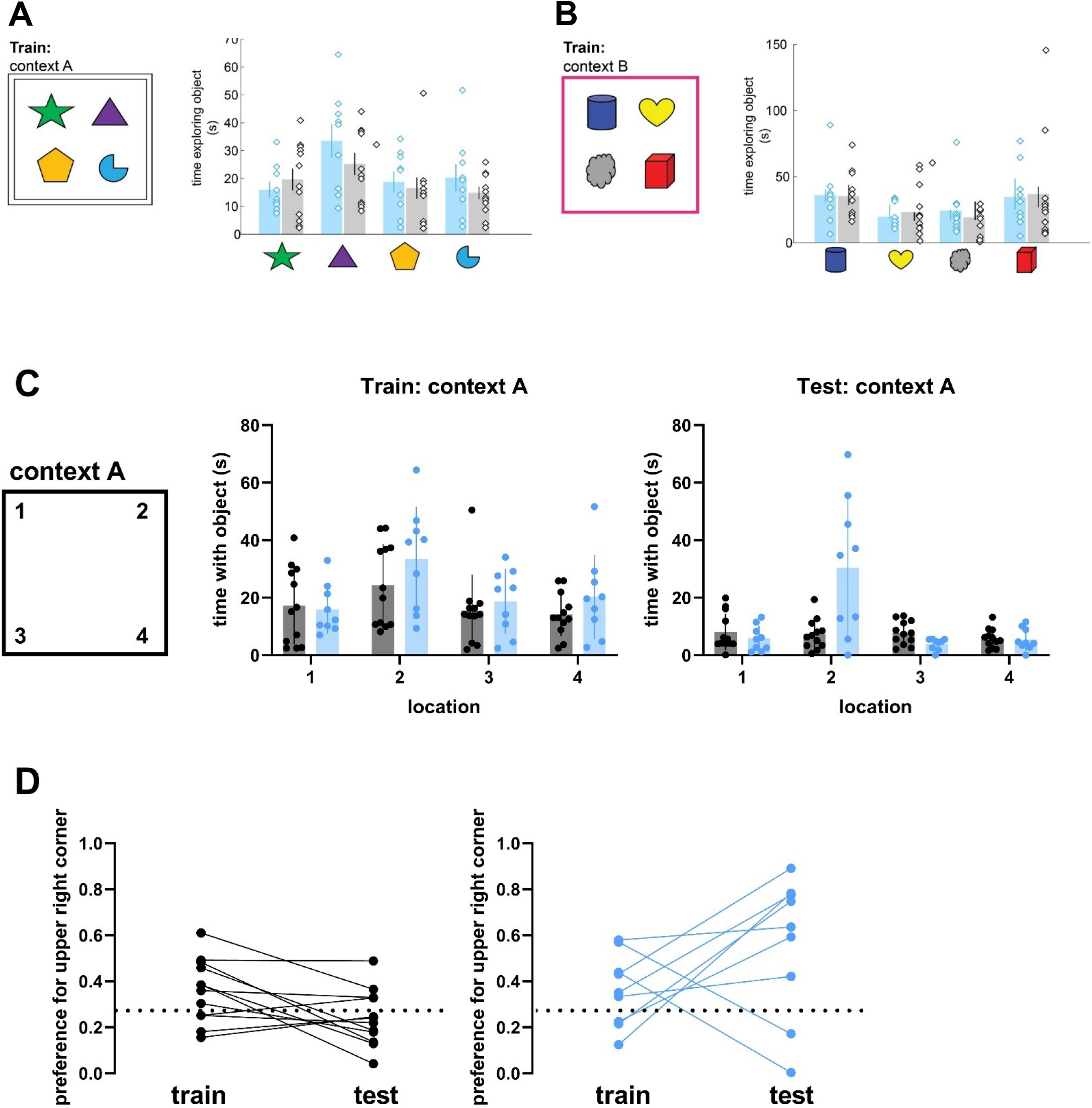
TeTx and control animals show no difference between groups in initial preference for objects. A) Cumulative time spent with each object in context A during the second training session (two-way repeated-measures ANOVA on one measure: A x B p-value = 0.56) in TeTx (blue) and control (black) animals. B) Cumulative time spent with each object in context B during the second training session (two-way repeated-measures ANOVA on one measure: A x B p-value = 0.94). C) Time spent with object in particular location during example training session and during testing. D) Preference for upper right corner (location 2) during second training session and testing in controls (black) and TeTx animals (blue). Preference is defined here as the amount of time spent in location 2/ total time spent in all 4 locations. Dashed line indicates 0.25, or no preference for location 2.

**Supplementary figure 3:**
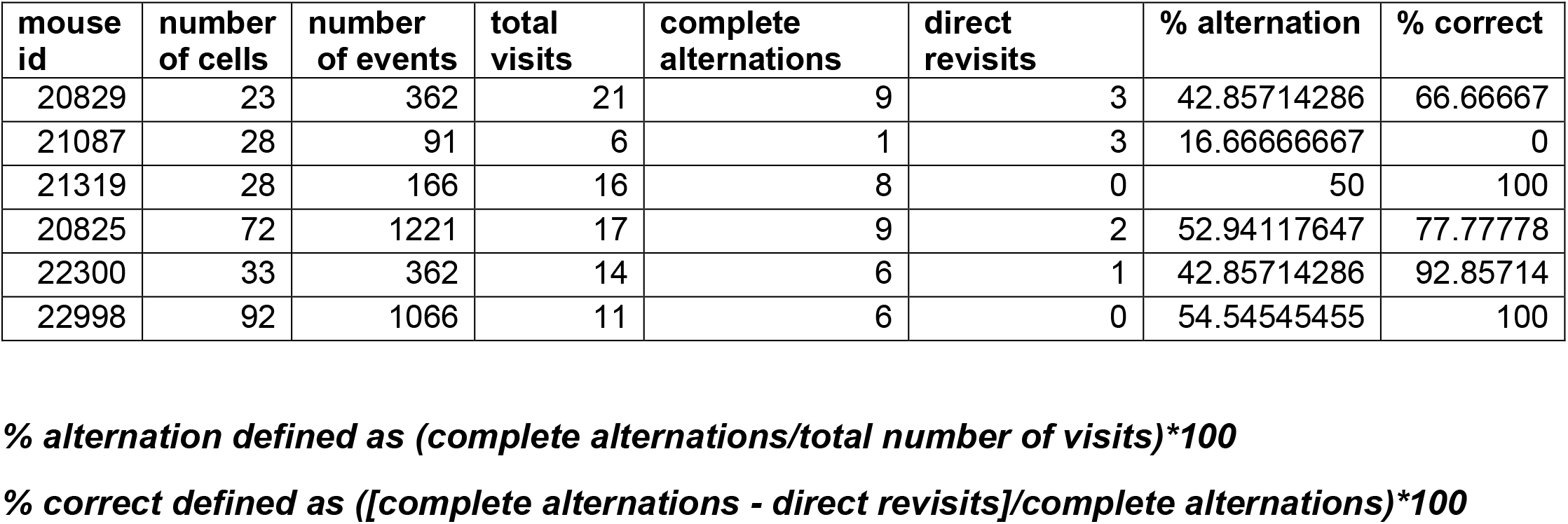
Additional behavioral data for Inscopix imaging. Table detailing the number of cells and calcium events imaged for each mouse, as well as the number of total visits, complete alternations, and direct visits. A complete alternation is a visit to each arm of the maze consecutively (e.g. arm A to B to C) while a direct revisit is a repeated visit to the same arm (e.g. arm A to arm A). The number of total visits includes the number of complete alternations, direct revisits, and indirect revisits (e.g. arm A to arm B to arm A).

## Methods

### Viral injections

For stereotaxic injections, adult mice were anesthetized through inhalation of 1.5% isofluorane gas and placed in a stereotaxic frame (Kopf Instruments). Mice were administered an analgesic (slow release buprenorphine, ~25 μL /25 g) subcutaneously before the beginning of surgery. After confirming that Lambda and Bregma were on the same dorsal-ventral plane, craniotomies were made with a high speed drill (Foredom MH-170) over prefrontal cortex (2.5 A/P, 1.0 L, 1.3 V) and/or lateral entorhinal cortex (−3.65 A/P, 4.5 L, 2.65 V and −4.15 A/P, 4.25 L, 2.15 V), in reference to the Allen Mouse Brain Atlas (Lein et al., 2007). Using a precision pressure injection system (Drummond Nanoject), a glass pipette filled with virus was lowered to the desired depth, briefly retracted (~0.2mm), and small amounts of virus were injected over a period of ~10 minutes (5-10 injections of 18-32 nL every 20 seconds). After waiting for an additional 5 minutes to prevent efflux of virus during pipette retraction, the glass pipette was retracted from the brain and the skin over the craniotomy was sutured shut. After applying several drops of topical anesthetic to the incision (bupivacaine), mice were allowed to recover under a heat lamp for 10-20 minutes and then placed in their home cage.

### Inactivation experiments

C57/BL6 mice (Jackson Laboratories) between the age of P60 and P120 (n = 12 experimental, 12 control) were injected with virally encoded Cre-dependent tetanus toxin or Cre-dependent GFP in PFC (~300 nl, AAV2/9 hSyn flex TeTx-YFP (made in house, plasmid courtesy of Fan Wang) or hSyn flex GFP, Addgene) and Cre in LEC (~140 nl at each site, AAVretro hSyn Cre, Addgene) according to the protocol described above. 4 weeks after viral injection, mice were transferred to the Duke Behavioral Core. One week after transfer, mice were tested in the Y maze and the Object context test. After behavioral testing, mice were perfused and histology was performed as described above. Only mice with bilateral viral expression contained within PL/IL and ORB across at least 3 consecutive coronal sections were included in this study. Two mice in the experimental cohort were excluded due to minimal (less than 3 consecutive sections containing PFC label) label in one of the hemispheres.

### Optogenetic experiments

C57/BL6 mice (Jackson Laboratories) between the age of P60 and P120 (n = 7 experimentals, 6 controls) were injected unilaterally with virally encoded Cre-dependent channelrhodopsin in PFC (~300 nl, AAV2/1 EF1a double floxed hChR2(H134R) mCherry-WPRE or flex hSyn GFP, Addgene) and Cre in LEC (140 nl at each site, AAVretro hSyn Cre, Addgene) according to the protocol described above. 4 weeks after viral injection, mice were implanted with a fiberoptic ferrule over the injection site (0.25 mm diameter, 0.3 NA, RWD) that was secured to the skull with dental cement (Metabond). 2 weeks after implantation, mice were transferred to the Duke Behavioral Core. 1 week later the mice were briefly anesthetized (isofluorane), coupled to a fiberoptic cable (ThorLabs) and laser (Shanghai Lasers, 493 nm) and allowed to fully recover for 10 minutes. Mice were then placed in the center of the Y maze and allowed to explore the maze for 10 minutes. Behavior was recorded in Ethovision XT and the laser was triggered every time the mouse entered the center of the maze (10 Hz pulse train for 1 second, PulsePal). After the experiment, the distance traveled, time spent in portions of the maze and the number of alternations were calculated in Ethovision. At the end of each experiment, mice were perfused and localization of viral expression and fiberoptic were visually confirmed with histology. Only mice with viral expression contained within PL/IL and ORB across 3 consecutive coronal sections and correct fiberoptic placement in prefrontal cortex were included in this study.

### One photon imaging in freely behaving mice

C57/BL6 mice (Jackson Laboratories) between the age of P60 and P120 (n = 6) were injected with virally encoded Cre-dependent GCaMP in PFC (~300 nl, AAV2/9 flex GCaMP6f, Addgene) and Cre in LEC (140 nl at each site, AAVretro hSyn Cre, Addgene) according to the protocol described above. Three weeks after AAV injection, a gradient index lens microendoscope (Proview Lens Probe, diameter 1.0 mm, length 4.0 mm, Inscopix) was implanted in the same position during a second surgery. Briefly, a small cranial window was made above the injection site in prefrontal cortex and the microendoscope was slowly lowered into the brain (100 um/minute to 1.7 mm DV). After 5 min of waiting, the lens was retracted back to 1.5 mm DV and fixed to the skull using adhesive luting cement (Metabond). A small screw was also affixed to the skull above the cerebellum to increase the stability of the implant. Cement, followed by a thin layer of tissue adhesive (VetBond) was used to seal the remaining exposed skull. One week after microendoscope implantation, a baseplate (Inscopix) was implanted over the lens, cemented in place and covered with a base plate cover (Inscopix).

5–6 weeks after initial viral injection, mice were habituated to the miniature microscope for 5–10 min before beginning the behavioral session. Calcium events were collected at 20 frames per second at 50% laser power (Inscopix nVista HD) as the animal explored a Y maze for 5 min. The onset of calcium imaging was synchronized with the placement of the animal in the arena for concurrent calcium imaging and behavioral tracking (Ethovision 11.5). At the end of each experiment, mice were perfused and localization of viral expression and lens placement were visually confirmed with histology. Only mice with viral expression contained within PL/IL and ORB across 3 consecutive coronal sections and correct lens placement in prefrontal cortex were included in this study.

The data collected from imaging sessions was decompressed (Inscopix Image Decompressor) and processed to extract calcium event timing (Inscopix Data Processing 1.2.1). Briefly, using the Inscopix software package mentioned above, data was spatially downsampled (2-3x) and motion-corrected in reference to the first frame of the recording. Cells were identified with PCA-ICA and confirmed as cells by eye. The calcium traces extracted from PCA-ICA were then thresholded (median absolute deviation: 4; event smallest decay time: 0.20 s) and the timing of calcium events from individual cells was calculated. The timing of events was used to calculate the rate of events for each cell (Matlab). The time at which the animal was in each portion of the maze was extracted from Ethovision XT and aligned with calcium transients (Matlab). Selectivity for the center of the Y maze was calculated as [(the number of events in the center of the maze/amount of time spent in center) – (number of events in the arms of the maze/amount of time spent in arms)] / [(the number of events in the center of the maze/amount of time spent in center) + (number of events in the arms of the maze/amount of time spent in arms)]. Center selective cells were defined as any cells with selectivity greater than 0. Arm selective cells were defined as any cells with selectivity less than 0.

### Behavioral paradigms and analysis

Mice were transferred to the Duke Behavioral Core facility at least 1 week before the beginning of behavioral experiments. One day before behavioral experiments, mice were moved to the room in which testing would occur in order to acclimate them to the testing room.

#### Y maze

We used the Y maze as a test of working memory in mice in which PFC-LEC cells were inactivated as well as in a separate cohort of mice in which PFC-LEC cells were optogenetically activated. A Y shaped maze with different visual cues on each arm was secured to the top of absorbent pads with tape on the floor of the testing room. Mice were placed in the center of the maze and allowed to spontaneously explore all arms of the maze for 5 minutes. Their behavior was recorded and the number of alternations as well as the time spent in each portion of the maze were analyzed in Ethovision XT. This task is unrewarded and has no delay period. Percent alternation was calculated as the actual number of correct alternations divided by the number of possible correct alternations.

#### Novel object-in-context

We used the novel object-in-context test as a measure of associative contextual memory in mice, following the protocol described by [13]. On day 1 (‘habituation’) mice were habituated to two different arenas containing no objects and distinguished by different high contrast patterns on the walls for 10 minutes in each context. On days 2-4 (‘training’), 4 unique objects were placed in the corners of each context and the mice were allowed to explore each context twice for 5 minutes separated by a 1-hour interval. The order of context exploration was counterbalanced on each training day. Critically, during training objects in context A are only encountered in context A and objects in context B are only encountered in context B. On day 5 (‘retention test’) two objects from A context were replaced by two objects from context B. Mice were then allowed to explore context A with the novel objects-in-context for 3 minutes. Mouse behavior was recorded and the distance traveled and the amount of time spent with each object was analyzed with Ethovision XT for all days of the paradigm. The discrimination index was calculated as (amount of time spent with novel objects-in-context – amount of time spent with familiar objects-in-context)/ total time spent with objects.

### Immunohistochemistry

Mice were deeply anesthetized (isofluorane) then perfused transcardially with phosphate buffered saline (PBS) followed by 4% paraformaldehyde in PBS (PFA). Brains were removed from the skull and placed in 30% sucrose PFA overnight for cryoprotection. Brains were then imbedded in OCT (Sakura) and frozen before sectioning (50 um sections, Cryostat). After placing free floating sections in wells of PBS, sections were either mounted on slides or stained for fluorescent markers. Sections that were stained were washed in 0.3% Triton X-PBS (PBS-T) solution (3 washes for 10 minutes each) then placed in blocking buffer (5% goat serum in PBS-T) for 2 hours at room temperature. Sections were then transferred to primary antibody solution (anti-Chick GFP, Sigma, 1:1000 in PBS-T; rabbit anti-glutaminase (Abcam #ab156876, 1:1000); chicken anti-GFP (Abcam #ab13970, 1:2000)) and placed on a shaker at 4 degrees overnight. After washing (3 washes in PBS-T for 10 minutes each), sections were placed in a secondary antibody solution (Alexa Fluor 568 goat-anti-rabbit (ThermoFisher #A11036, 1:1000); Alexa Fluor 488 goat-anti-chicken (Jackson ImmunoResearch #103-545-155, 1:2000)) for 1 hour at room temperature then washed again and mounted on slides. Images were then acquired on a Zeiss Apotome (tile scans) or a Zeiss 710 (z-stacks).

## Notes

### Competing Interest Statement

The authors have declared no competing interest.

## References

1. D’Esposito, M. and B.R. Postle, The cognitive neuroscience of working memory. Annu Rev Psychol, 2015. 66: p. 115–42.

2. Josselyn, S.A. and S. Tonegawa, Memory engrams: Recalling the past and imagining the future. Science, 2020. 367(6473).

3. Witter, M.P., et al., Architecture of the Entorhinal Cortex A Review of Entorhinal Anatomy in Rodents with Some Comparative Notes. Front Syst Neurosci, 2017. 11: p. 46.

4. Montchal, M.E., Z.M. Reagh, and M.A. Yassa, Precise temporal memories are supported by the lateral entorhinal cortex in humans. Nat Neurosci, 2019. 22(2): p. 284–288.

5. Keene, C.S., et al., Complementary Functional Organization of Neuronal Activity Patterns in the Perirhinal, Lateral Entorhinal, and Medial Entorhinal Cortices. J Neurosci, 2016. 36(13): p. 3660–75.

6. Schultz, H., T. Sommer, and J. Peters, The Role of the Human Entorhinal Cortex in a Representational Account of Memory. Front Hum Neurosci, 2015. 9: p. 628.

7. Nilssen, E.S., et al., Neurons and networks in the entorhinal cortex: A reappraisal of the lateral and medial entorhinal subdivisions mediating parallel cortical pathways. Hippocampus, 2019. 29(12): p. 1238–1254.

8. Zhang, Y., et al., Identifying local and descending inputs for primary sensory neurons. J Clin Invest, 2015. 125(10): p. 3782–94.

9. Tervo, D.G., et al., A Designer AAV Variant Permits Efficient Retrograde Access to Projection Neurons. Neuron, 2016. 92(2): p. 372–382.

10. Chao, O.Y., et al., The medial prefrontal cortex-lateral entorhinal cortex circuit is essential for episodic-like memory and associative object-recognition. Hippocampus, 2016. 26(5): p. 633–45.

11. Wilson, D.I., et al., Lateral entorhinal cortex is critical for novel object-context recognition. Hippocampus, 2013. 23(5): p. 352–66.

12. Wilson, D.I., et al., Lateral entorhinal cortex is necessary for associative but not nonassociative recognition memory. Hippocampus, 2013. 23(12): p. 1280–90.

13. Lesburgueres, E., et al., The Object Context-place-location Paradigm for Testing Spatial Memory in Mice. Bio Protoc, 2017. 7(8).

14. Chen, T.W., et al., Ultrasensitive fluorescent proteins for imaging neuronal activity. Nature, 2013. 499(7458): p. 295–300.

15. Kraeuter, A.K., P.C. Guest, and Z. Sarnyai, The Y-Maze for Assessment of Spatial Working and Reference Memory in Mice. Methods Mol Biol, 2019. 1916: p. 105–111.

16. Riley, M.R. and C. Constantinidis, Role of Prefrontal Persistent Activity in Working Memory. Front Syst Neurosci, 2015. 9: p. 181.

17. Lee, J.H., et al., Global and local fMRI signals driven by neurons defined optogenetically by type and wiring. Nature, 2010. 465(7299): p. 788–92.

18. Callaway, E.M. and L. Luo, Monosynaptic Circuit Tracing with Glycoprotein-Deleted Rabies Viruses. J Neurosci, 2015. 35(24): p. 8979–85.

19. Gilmartin, M.R., et al., Prefrontal activity links nonoverlapping events in memory. J Neurosci, 2013. 33(26): p. 10910–4.

20. DeNardo, L.A., et al., Temporal evolution of cortical ensembles promoting remote memory retrieval. Nat Neurosci, 2019. 22(3): p. 460–469.

21. Grossberg, S., Desirability, availability, credit assignment, category learning, and attention: Cognitive-emotional and working memory dynamics of orbitofrontal, ventrolateral, and dorsolateral prefrontal cortices. Brain Neurosci Adv, 2018. 2: p. 2398212818772179.

22. van Strien, N.M., N.L. Cappaert, and M.P. Witter, The anatomy of memory: an interactive overview of the parahippocampal-hippocampal network. Nat Rev Neurosci, 2009. 10(4): p. 272–82.

23. Fernandez-Ruiz, A., et al., Gamma rhythm communication between entorhinal cortex and dentate gyrus neuronal assemblies. Science, 2021. 372(6537).

24. Anderson, M.C., J.G. Bunce, and H. Barbas, Prefrontal-hippocampal pathways underlying inhibitory control over memory. Neurobiol Learn Mem, 2016. 134 Pt A: p. 145–161.

25. Davis, R.L. and Y. Zhong, The Biology of Forgetting-A Perspective. Neuron, 2017. 95(3): p. 490–503.

